# Active sensing in bees through antennal movements is independent of odor molecule

**DOI:** 10.1101/2021.09.13.460114

**Authors:** Nicolas Claverie, Pierrick Buvat, Jérôme Casas

## Abstract

When sampling odors, many insects are moving their antennae in a complex but repeatable fashion. Previous works with bees have tracked antennal movements in only two dimensions, with a low sampling rate and with relatively few odorants. A detailed characterization of the multimodal antennal movement patterns as function of olfactory stimuli is thus wanting. The aim of this study is to test for a relationship between the scanning movements and the properties of the odor molecule.

We tracked several key locations on the antennae of 21 bumblebees at high frequency (up to 1200 fps) and in three dimensions while submitting them to puffs of 11 common odorants released in a low-speed continuous flow. To cover the range of diffusivity and molecule size of most odors sampled by bees, compounds as different as butanol and farnesene were chosen, with variations of 200% in molar masses. Water and paraffin were used as negative controls. Movement analysis was done on the tip, the scape and the base of the antennae tracked with the neural network Deeplabcut.

Bees use a stereotypical motion of their antennae when smelling odors, similar across all bees, independently of the identity of the odors and hence their diffusivity. The variability in the movement amplitude among odors is as large as between individuals. The first oscillation mode at low frequencies and large amplitude (ca. 1-3 Hz, ca. 100°) is triggered by the presence of an odor and is in line with previous work, as is the speed of movement. The second oscillation mode at higher frequencies and smaller amplitude (40 Hz, ca. 0.1°) is constantly present. Antennae are quickly deployed when a stimulus is perceived, decorrelate their movement trajectories rapidly and oscillate vertically with a large amplitude and laterally with a smaller one. The cone of air space thus sampled was identified through the 3D understanding of the motion patterns.

The amplitude and speed of antennal scanning movements seem to be function of the internal state of the animal, rather than determined by the odorant. Still, bees display an active olfaction strategy. First, they deploy their antennae when perceiving an odor rather than let them passively encounter it. Second, fast vertical scanning movements further increase the flow speed experienced by an antenna and hence the odorant capture rate. Finally, lateral movements might enhance the likelihood to locate the source of odor, similarly to the lateral scanning movement of insects at odor plume boundaries. Definitive proofs of this function will require the simultaneous 3D recordings of antennal movements with both the air flow and odor fields.

## 1 Introduction

Active sensing can be characterized as the spending of energy to probe the environment. Biological examples include echolocation in bats or electrolocation in fishes. Contrary to passive sensing, active sensing allows the animal to gather additional information on its environment by varying its sensing behavior (^1^). Active sensing behavior can be found in the context of olfaction in mammal sniffing (even underwater ^2^) were the sniffs frequency (^3,4^) and nose motion (^5^) can be varied to improve sensing, and where the expiration jets are used to increase the capture of odorant (up to 18 times in Staymates et al ^6^). Crustaceans have also been observed using jets to improve their odorant capture (^7,8^). Active sensing in insect includes casting behavior in flying insects (^9–12^) and larvae (^13^) that improve odorant source tracking, and insect wing motion that allow improved sampling (^14–17^) with increase in capture rate by the antennae that can reach more than an order of magnitude compared to still air (^18^).

In Crustaceans, antennae are the main organs of olfaction (^19^), and antennal motion as an active sensing mechanism is well described (^20–23^). The motion of the antennules increases the water flow on the aesthetascs, increasing thereby their leakiness and allowing molecules to contact the receptors much easier. In insects, the antennae are not only an olfaction organ, but function as multisensory organs too (^24,25^). Their motion in the context of flight (^26–28^) and mechanoreception (^29^) have been well studied, especially for stick insects where the active tactile sensing is quite present (^30–34^). The role of antennal motion in olfaction is less understood. In cockroaches, variations of the antennae position and motion was observed to be depending on the stimulus (^35^) and odor plume structure (^36^). In locusts, the antennal motion was linked to increased intermittency of the odor signal that could help stimulus localization (^37^). Those insects have antennae with short scape, so their motion is mostly limited to rotations around their base. Bees and ants, due to their longer scape, have antenna capable of more complex movement. Ants antennal motion while tracking a scent trail was recently characterized by Draft et al (^38^). They discovered several distinct stereotyped patterns of antennal motion used by ants following a trail.

Bees do not use their antennae to trace scent trails in the way ants do. Previous studies on the characteristics of bee’s antennae motion are scarce and have tracked antennal movements in only two dimensions, with a low temporal sampling rate and with relatively few odorants (^39–44^). Some of their results were contradictory, with Erber et al (^40^) and Suzuki (^39^) reporting that antennae moved toward the source of odor whereas Chole et al (^43^) and Birgiolas et al (^44^) reporting the antenna as moving away from the odor before conditioning. Lambin et al (^42^) reported an increase in the antennae angular speed but Chole et al (^43^) noticed no increase in speed for unconditioned stimuli. Peteraitis (^41^) was able to show that an increase in odorant flow speed on the antennae comparable to the stroke speed of antennal motion increased the EAG response of workers honeybee. This paper is to our knowledge the only one linking bees’ antennal motion to increase in odorant capture, and the only one to propose a mechanism for the role of antennal motion in the olfactory process of bees.

Therefore, the use of active sensing by bees to smell odors remains an open question. A detailed characterization and a three-dimensional analysis of the multimodal antennal movement patterns as function of olfactory stimuli is thus lacking. This is our aim. Our hypothesis is that the bees’ antennal movement sample the air around the insect in a way that improve odorant capture. Such an active sampling behavior of the insect might be adapted depending on the chemical characteristics of the stimulus. Either because the distribution of the odor in the air is varying or because its capture dynamic depends on those properties. Both should be affected by the diffusivity and the molar mass of the molecule. The goal of this study was then to measure the antennal motion of bumblebees and their variability for different compounds.

Our analysis was done of bumblebees, due to their large size making observations easier, and due to the ease of acquiring individuals. Bumblebee present similar antenna structure and sensilla types to other bees of the Apidae family (^45^) so we assume that their antennal motion would be representative of bees in general. As stimuli, we chose a panel of pure plant volatile compounds with variable diffusivity ranging from 4.9 to 9.1 mm^2^/s and molecular weight ranging from 74 to 204 g/mol, covering the common range of values for plant volatile compounds encountered by bees (^46^). The diffusivity of the molecules was calculated using the method of Fuller et al (^47^). All the odor we used had been used before in bees olfactory testing, to ensure that bumblebees can smell them. Using pure odorant ensured that we could test the effect of chemical characteristic of the molecules. Since our bees were free to forage outside, they were exposed to blends of plant odors but should not have developed an attractive or repulsive response to any of our pure stimuli (^48^).

## 2 Material and methods

### 2.1 Experimental design

We used a commercial hive (Standard Hive (B.t.), Biobest Group NV, Westerlo, Belgium) of bumblebees (*Bombus terrestris*) for our experiments. This type of hive contains a queen and around 80 workers at delivery. It also included food for the colony. A hive was installed outside of the laboratory in Tours, France according to the company recommendation. The bumblebees were free to forage in the environment. All tested bumblebees were captured when they exited the hive to forage. Once or twice a day, a worker bumblebee was captured and put in a plastic container. In order to conduct olfactory tests, it was then transferred into a 10mL plastic tube with the end cut off, to allow for the head to pop out, but not the rest of its body (inspired by Ma et al ^49^) (Fig 1A). To force the bumblebee to keep its head outside, a piston with a conical cavity in its center designed to accommodate the insect abdomen pushed the bee to the end of the tube. The method to place the insect inside the fixation tube involved guiding the insect from the plastic container into smaller and smaller tubings until it entered the fixation tube by itself. This method was chosen to avoid using CO_2_ or cold to anesthetize the insect.

**Fig 1:**
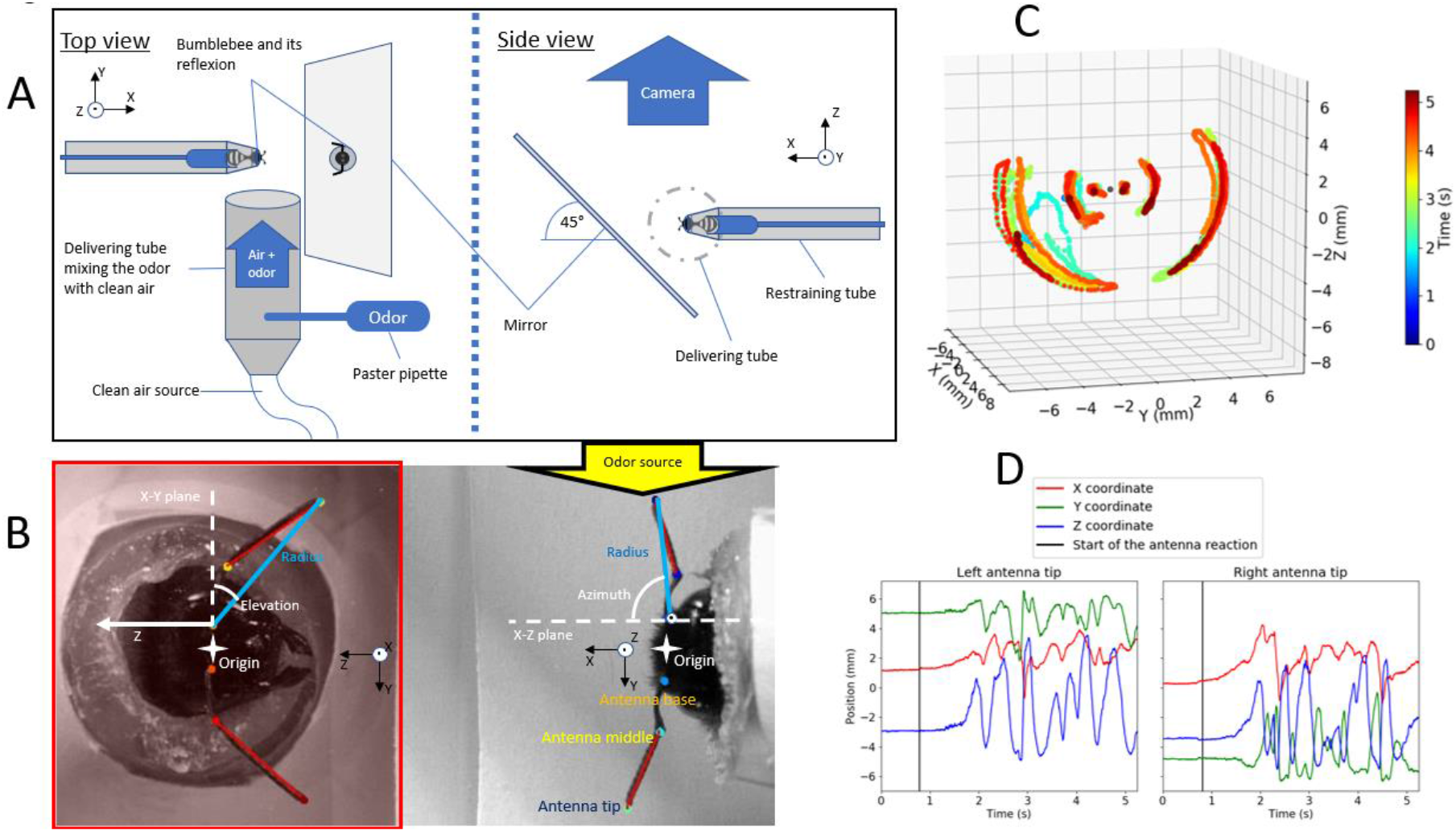
Experimental set-up and typical data results. (A) Schematics of the experimental setup for restraining, stimulating, and filming the bumblebees (top and side view). (B) Image captured by the camera and marked by the tracking software. The coordinates system (spherical system) used for the analysis is also described. (C) Example of the data extracted from the video. Here a 3D reconstruction of the trajectory of the tip the middle and the base of each antenna during a test. The color of the curve represent time. (D) Evolution of the x y and z coordinates (in mm) of the tip of the antenna over time with respect to the head center.

The fixation tube was then placed in front of a mirror inclined at 45 degrees and below a high-speed camera. The mirror was used to provide a second point of view and to measure the altitude of each point of the antennae, while the other coordinates were extracted from the direct image of the insect (Fig 1B). This allowed a 3D reconstruction of the antennae motion. The camera (Phantom V9; Vision Research, Perth, Australia) recorded, in the same frame, both the top and the front views of the insect through the mirror. It recorded at 300 fps for 5 seconds, with around 0.045 mm/pixel resolution and in grayscale visible light for the standard trial. To better capture fast and small-scale motion of the antennae, zoomed measurement were also recorded at 1200 fps during 1.5 second with 0.001mm/pixels resolution. The stimuli were delivered from the right side of the insect through a plastic tube with a diameter of 27mm, called thereafter the “delivering tube”. The delivering tube is made using a 50mL plastic test tube with a conical base. A hole was drilled at the bottom of the base to allow for a pipe of 1cm in diameter. A pump was delivering a constant flux of air through that pipe, acting as a carrier for the stimulus. The stimulus was delivered by a plastic Pasteur pipette through a small hole on the side of the delivering tube. The odor from the pipette and the air from the pump then mixed and travelled inside the delivering tube over its length (8cm) and exited next to the head of the insect (Fig 1A). A quantitative flow tracing was done by filming the progression of DEHS (di2-ethylhexyl sebacate) droplet mist inserted in the tube in the same way the odor was. It showed that the airflow was laminar with an average speed around 35mm/s (±5mm/s) measured by manually tracking the particles in the videos of five different trials. Despite the odor concentration not being homogeneously mixed, it was well distributed inside the airflow, ensuring a continuous contact with some portion of the insect antennae (Fig S1).

### 2.2 Odor preparation

Common plant odors (^50–52^) of varying molecular weights and diffusion coefficients were selected as stimuli for the experiment. Our goal was to evaluate the suspected effect of the molecular weight and the diffusivity on the motion of the antenna. We chose common plant odors with large differences in diffusivity, like butanol (9.1 mm^2^/s) and farnesene (4.9 mm^2^/s). We also tested molecules with similar properties, like ocimene and alpha pinene that are structural isomers but with very different scents. The diffusion coefficients of the molecules used were calculated using the method of Fuller et al (1966) as adapted by Tang et al (2015). The odors used were benzaldehyde (≥99%, Sigma-Aldrich, Taufkirchen, Germany), 1-butanol (99%, Alfa Aesar, USA), phenylacetaldehyde (95%, Alfa Aesar), + alpha pinene (analytical standard, Fluka, Germany), nonanal (≥95% Sigma-Aldrich), farnesene (mixture of isomers, Sigma-Aldrich), beta caryophyllene (≥80%, Sigma-Aldrich), R+ limonene (analytical standard, Fluka), trans-2-hexen-al (≥95%, Sigma-Aldrich), trans-2-hexen-ol (≥95%, Sigma-Aldrich) and ocimene (≥90%, Sigma-Aldrich). All those odors can be sensed by bees (^53–61^) (Table 1). All odors were diluted in water except for phenylacetaldehyde that had to be diluted in paraffin oil (puriss, Sigma-Aldrich) for solubility reasons. The dilution ensured that the insect would not be overwhelmed by the intensity of the odor and would display normal antennal motion to better detect stimuli. The dilution rate was 200 times for all odors, excepted for alpha pinene and butanol that needed to be less diluted (20 times) to elicit a response. The diluted mixture was then used to rinse the inside of a plastic Pasteur pipette for a short time. The adsorption of the odor by the pipette’s walls was sufficient to deliver stimuli when pressing the pipette. In total, 14 different stimuli could be given to each insect, namely 11 odors plus water, paraffin oil and ambient air as controls. To avoid cross-contamination, several pipettes and delivering tubes (the tube mixing the odor from the pipette with the air) were used, one for each odor plus one for the controls. Each stimulus was delivered to bumblebees in a randomized sequence.

**Table 1:**
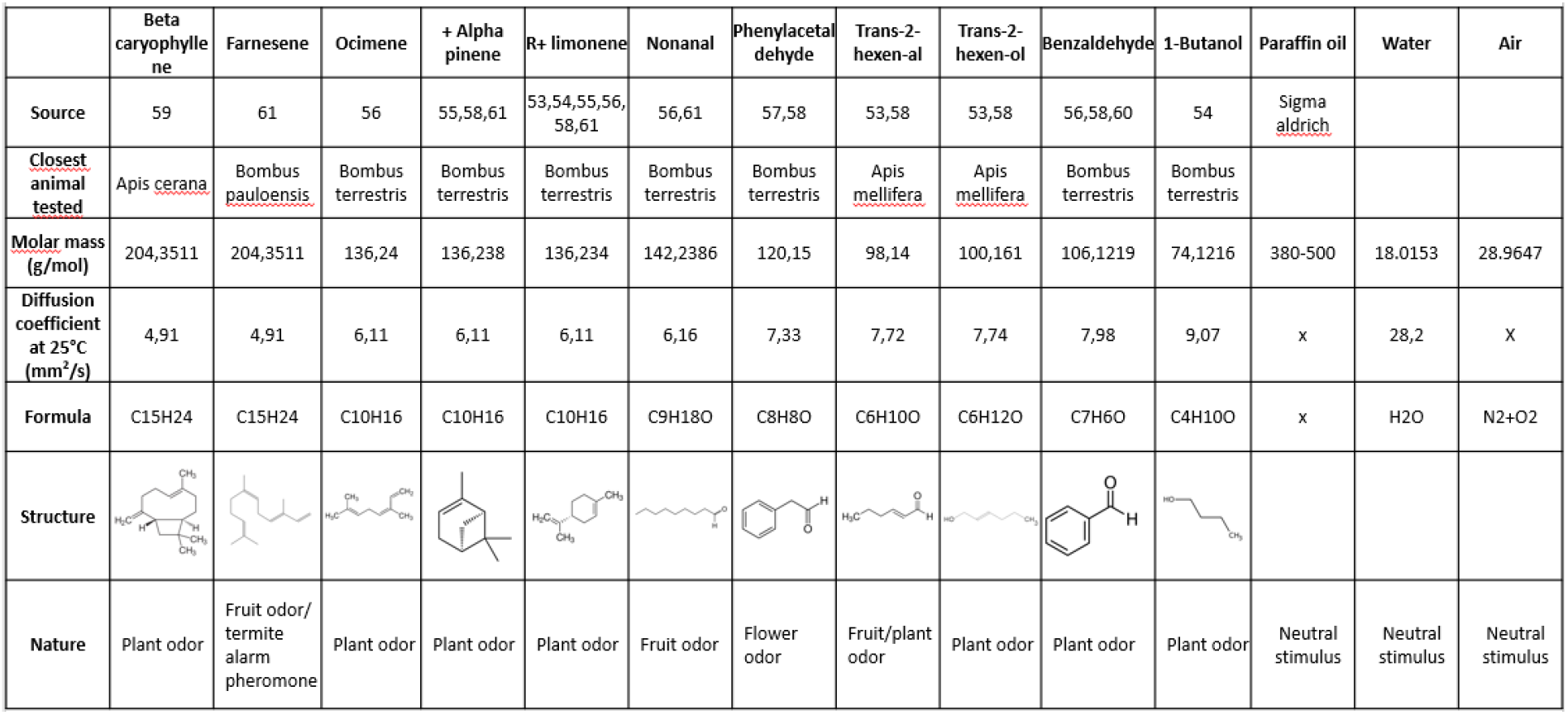
Odor used, their physical properties, studies that previously used them for odor testing and the insect closest to the *Bombus terrestris* they were tested on.

|  | Beta caryophyllene | Farnesene | Ocimene | + Alpha pinene | R+ limonene | Nonanal | Phenylacetaldehyde | Trans-2-hexenal | Trans-2-hexenol | Benzaldehyde | 1-Butanol | Paraffin oil | Water | Air |
| --- | --- | --- | --- | --- | --- | --- | --- | --- | --- | --- | --- | --- | --- | --- |
| Source | 59 | 61 | 56 | 55,58,61 | 53,54,55,56,58,61 | 56,61 | 57,58 | 53,58 | 53,58 | 56,58,60 | 54 | Sigma aldrich |  |  |
| Closest animal tested | *Apis cerana* | *Bombus pauloensis* | *Bombus terrestris* | *Bombus terrestris* | *Bombus terrestris* | *Bombus terrestris* | *Bombus terrestris* | *Apis mellifera* | *Apis mellifera* | *Bombus terrestris* | *Bombus terrestris* |  |  |  |
| Molar mass (g/mol) | 204,3511 | 204,3511 | 136,24 | 136,238 | 136,234 | 142,2386 | 120,15 | 98,14 | 100,161 | 106,1219 | 74,1216 | 380-500 | 18.0153 | 28.9647 |
| Diffusion coefficient at 25°C (mm<sup>2</sup>/s) | 4,91 | 4,91 | 6,11 | 6,11 | 6,11 | 6,16 | 7,33 | 7,72 | 7,74 | 7,98 | 9,07 | x | 28,2 | X |
| Formula | C15H24 | C15H24 | C10H16 | C10H16 | C10H16 | C9H18O | C8H8O | C6H10O | C6H12O | C7H6O | C4H10O | x | H2O | N2+O2 |
| Structure |  |  |  |  |  |  |  |  |  |  |  |  |  |  |
| Nature | Plant odor | Fruit odor/termite alarm pheromone | Plant odor | Plant odor | Plant odor | Fruit odor | Flower odor | Fruit/plant odor | Plant odor | Plant odor | Plant odor | Neutral stimulus | Neutral stimulus | Neutral stimulus |

### 2.3 Experiment procedure

After being fixated, bumblebees are usually very active and move their antennae erratically. In order to study the reaction of an insect in a calm state, it was put in the dark at ambient temperature until its antennae stopped moving. The insect was then carefully affixed in front of the mirror in the recording zone. The pump was turned on, so that clean air started to blow gently on the insect head trough the delivering tube at a speed of 35mm/s. If the insect remained calm, the pipette was affixed to the delivering tube and pressed three times fully, within about one second. About three second later, the camera post-trigger was pressed to record the last five seconds of the video. This allowed for the camera to capture one or two seconds of the insect movement before the stimulus delivery, the onset of the scanning behavior and two or three seconds after the detection of the stimulus. The insect was then placed back in the dark to calm down, the delivering tube was changed for the next stimulus and the video was saved to the computer. Each stimulus was separated by at least 10 minutes. Once all stimuli were delivered or once the bumblebee stopped responding to any stimulus, the insect was marked with acrylic paint and released next to the hive. No bumblebees were reused in following trials thanks to their markings. Most bumblebees became tired before receiving the last stimulus. This meant that in the end, even if 21 different worker bumblebees were tested, they did not receive every odor, and so each odor was only tested on 7 to 15 bumblebees.

Additionally, videos of excited bumblebees in the absence of odors were recorded for comparison. Zoomed videos were also recorded to look for small scale and high-speed movements of the antennae, both with and without stimulus. The constrains of the higher framerate and magnification made it impossible to capture the full onset of the reaction and to track the large oscillation of the antennae accurately. This was due to the zoomed videos being only 1 second long and having a narrow depth of field.

In total, 24 different bumblebees were tested with this protocol but 2 were discarded after injuries of the antennae were found on the videos and 1 was discarded for having never responded to a stimulus. 225 videos were used for analysis.

### 2.4 Tracking of the antennae

The tracking of the antennae was done using the deep neural network software DeepLabCut, a software package mostly used for animal pose estimation (^62,63^). The videos were first compressed, cropped to be as tight as possible on the bumblebee, while ensuring that the antennae were always within frame, and saved as AVI files using ImageJ^64^. Then, the videos were all loaded inside DeepLabCut. To train the algorithm, 226 frames were extracted and annotated manually from 44 different videos. The points annotated were the tip of the flagellum, the base of the antenna and the thickest part of the pedicel, on both antennae and their reflections in the mirror (Fig 1B). The frames were manually chosen to be as different as possible from each other and to cover the range of antennal motions. The architecture of the neural network used was the Resnet 152 (^65^). Other Resnet and mobilenet type networks were tested but gave unsatisfactory results. The default augmentation method of the software was used (the software creates slightly modified copies of the annotated image to artificially increase the training sample of the neural network). The network was then trained for 69000 iterations using the default training step sizes until the error of the network reached a plateau. The final positioning error on the testing set was around 4.7 pixels which was low enough for our application since it is of the order of the diameter of the antenna. The network was then applied to all videos.

### 2.5 Data analysis

The neural network returned a csv file for each video containing the x and y coordinates, in pixels, of the tip of the flagellum, the pedicel, and the base of the antenna for both antennae and their reflected image in the mirror for a total of 12 points per frames. For each point, the network also recorded its confidence in the position of the point for each frame. The videos were around 1800 frames long. For the “zoomed” videos, their field of view was too small to have both the top view and the mirror image. So, the coordinates were recorded only in 2D with only 6 points per frames (Fig 1B).

In order to analyze the obtained data, each csv file was loaded in python 3.7.4^66^ using the package csv. The point between the mean position of the base of each antenna was chosen as the origin (the mirror image of the base of the antennae provided the height coordinate z). A coordinates system was then created, centered on the origin. The position of each point of the antennae was expressed in this system (Fig 1B). The known size of the delivering tube on the camera was used to convert the coordinate from pixels to millimeters. The time of each frame was obtained by scaling the frame number by the frame per second setting of the camera. The average video length was 6 seconds for the 300 fps videos and 1.5 seconds for the “zoomed” videos. The length and diameter of a typical antenna was also recovered from the video.

All plots were realized using the matplotlib.pyplot 3.1.1 library^67^. Using the obtained coordinates, a 3D plot of the trajectory of each point of the antenna and of the origin was realized (Fig 1C). This revealed that the motion of each antenna is very similar to a rotation around the base point of the antenna. Because of this, the antennae point coordinates were converted from a cartesian to a spherical referential with the radius (mm), azimuth (°), and elevation (°) as coordinates (Fig 1B) (see eq 1).

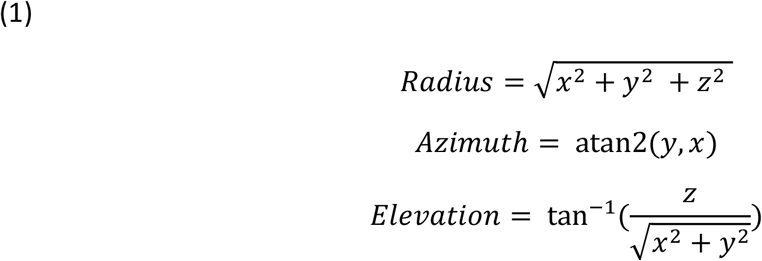

With x, y and z being the cartesian coordinates of the point in mm. The average position of base of the corresponding antenna is used as the origin. Using the fact that antennae do not flex much, the motion of the tip was described in a spherical coordinate system using two angles and one length per antenna. That way, the length coordinate called radius measures the extension of the antenna (the radial component of the motion) and the two angles describe its orientation, or angular position. The elevation angle represents the position in the vertical plane and the azimuth angle the position in the horizontal plane (Fig 2).

**Fig 2:**
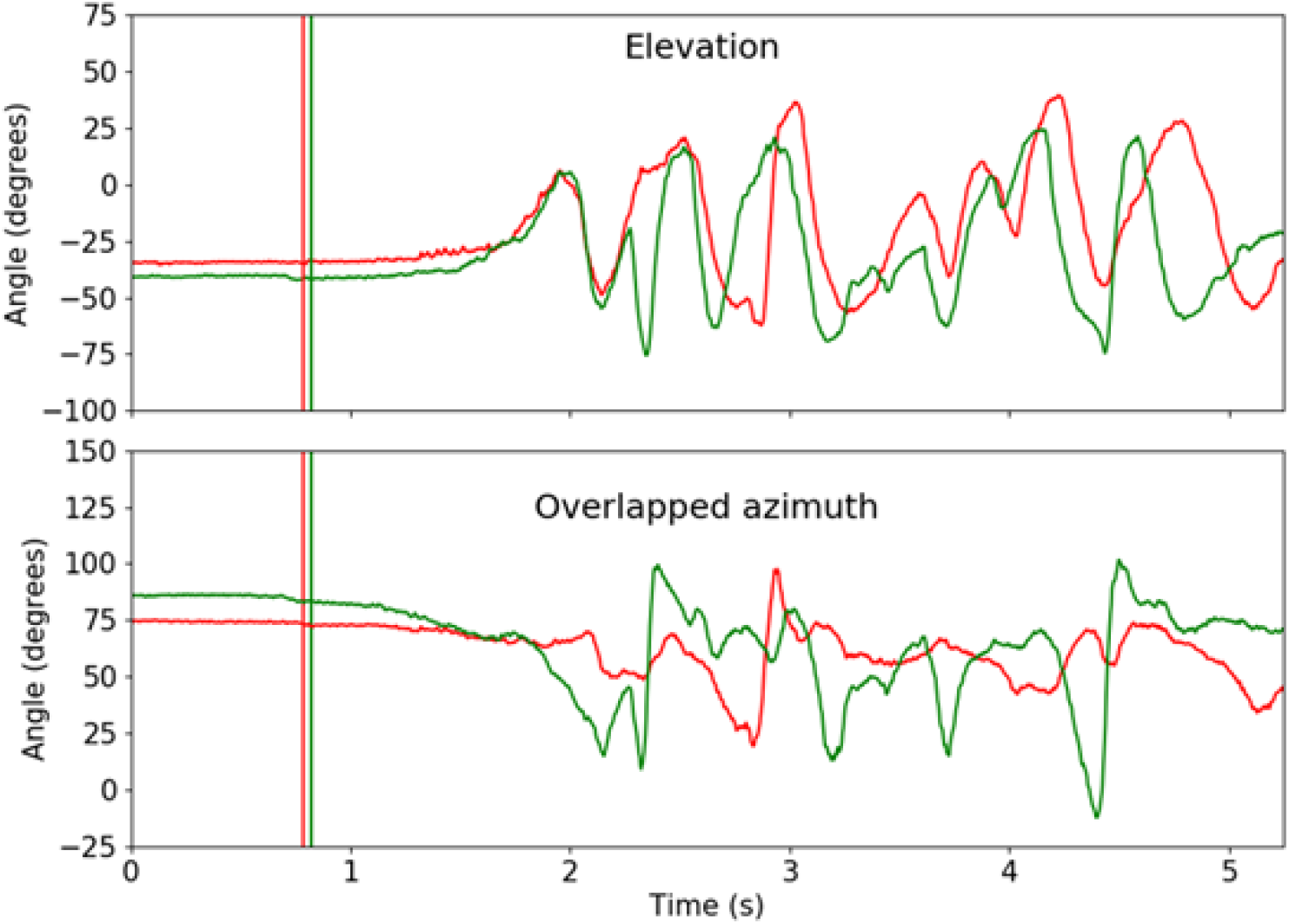
Example of superposition of the angular position of the tip of both antennae with respect to time. In green the right antenna and in red the left antenna. The sign of the right azimuth was inversed to allow for the overlapping of the curves. Each vertical bar marks the start of the reaction of one antenna (the green bar for the right antenna and the red for the left).

To pinpoint the start of the reaction of a bumblebee to an odor stimulus, a baseline is created using the position of the tip of the antenna during the first second of the video. This gives a mean and a standard deviation for the antenna position of this resting bumblebee. The start of the reaction is then determined as the last frame where the position of the tip of the antenna is within one standard deviation of the baseline and then increases gradually to above five standard deviations, without coming back within the threshold.

To evaluate the speed of the motion of each marker in the video, the difference in their 3D position between two frames was measured and divided by the time between those frames. To limit the impact of the tracking noise, we measured the displacement of frames separated by 1/60th of a second instead of consecutive frames because it reduced the noise, and the main frequencies of the motion were way below 30Hz. The speed of the high frequency oscillation was analyzed in the zoomed videos at 1200 fps.

As a primary filter against mislabeled points, if a detected point was too far from the insect to belong to the antennae, the point position was discarded and its position from the previous frame was kept. The maximum distance of a point from the origin of the coordinate was 10 mm in the Y direction, 8 mm in the Z direction, or 8 mm in front or 3 mm behind the insect. The confidence in the point position given by the neural network was also used to evaluate the quality of the entire video sequence. Each frame with at least one marker confidence below 75% was counted as a “bad” frame. The quality of a video was then calculated as one minus the ratio of “bad” frames (with at least one uncertain marker) over the total number of frames. For example, if 20 percent of the frame had at least one maker confidence below 0.75, the quality of the video would be 0.8. Those bad frames were also removed from the evaluation of values that were sensitive to extreme values, such as the maximum amplitude of the antenna motion or the maximum stroke speed. If the quality of the video was below 0.5, the video was discarded to avoid tracking error polluting the analysis. This left 149 good quality videos, of which 121 are response to an odor, that were used in the following analysis.

From these data, we were able to plot the evolution of the cartesian and polar coordinates of the tip of the antennae (Fig 1D, Fig 2), plot the trajectory of each point in 3D (Fig 1C), plot the volume of air sampled by each antenna and the speed of each antenna with respect to time. We could also mark the starting point of the reaction of each antenna. To extract the frequency content of the temporal movement of the antennae, and eventually identify a particular pattern of oscillation, we performed a spectral analysis using fast Fourier transforms of the temporal evolution of the angular coordinates of each antenna using the python package scipy.fftpack of Scipy 1.6.1 (^68,69^). In order to get an approximation of the volume of air sensed by an antenna, its size and diameter was measured to calculate the lengthwise cross-sectional area of the antenna. By multiplying this value by the instantaneous displacement of the antenna in the airflow, the volume of air crossed by the antenna was calculated for each frame and then summed to obtain the total volume sampled over time. To attenuate the effect of the positioning noise, the displacement of the antenna was calculated over frames separated by 1/60th of a second. We took into account the motion of the air around the antenna due to the applied airflow which caused a passive sampling rate.

## 3 Results

### 3.1 General patterns of motion

The general pattern of antennal movements is the same for all bees. At the start of the stimulation, the antennae are mostly immobile since the insect is calm, except for some very small and fast oscillations. Once the antennae start reacting to the stimulus, they deploy abruptly (the antennae go from nearly static to their full amplitude of motion in less than 0.5 seconds) and begin a stereotypical movement made of a combination of up and down and left and right slow oscillations. This scanning behavior is similar for all tested bees but can vary in amplitude, frequency, and overall regularity. In parallel to those slow oscillations, the fast oscillations of the antenna flagellum are still present, but their amplitude is very small compared to one of the slow oscillations. Those fast vibrations seem to occur all the time, even without stimuli. Zoomed videos had to be recorded to characterize them precisely. Plotting the trajectory of the markers in 3D shows that, during its motion, an antenna keeps a mostly fixed flexion angle between flagellum and scape (Fig S2). The antenna flexes by plus or minus 10 degrees from a mean position, which corresponds to a radial motion at the tip of around 5% of the length of the antennae. Therefore, the dominating movement is a rotation of the whole antenna, not flexion.

To evaluate the global pattern of antennal angular motion, we overlapped the angular positions they visited in a matrix plot summing all trials. The antennae move in the same angular region around the insect head. We can see two separated regions, one for each antenna. The antennae move at an azimuth of 20 to 80 degrees and an elevation of -75 to 50 degrees for all stimuli and bumblebee (Fig 3). This matrix plot is showing the most frequently visited locations of the antennae. To evaluate the dynamical nature of the motion, the distribution of the instantaneous direction of motion (the direction in which the tip of the antenna moves from one frame to the next) of the antenna tip was also examined for all recordings. Compilations were made for each insect and each molecule. We can see that the antennae move in specific directions (Fig S3). The most prevalent direction of motion and the direction of largest motion is near the vertical (90° and 270° directions) for both antennae with the quickest motion in the downward direction. This mean that the instantaneous antennae motion is mostly an up and down oscillation, with a slight inclination of this direction between zero and 45 degrees from the vertical, such that their joint motion forms a V shape (Fig S3).

**Fig 3:**
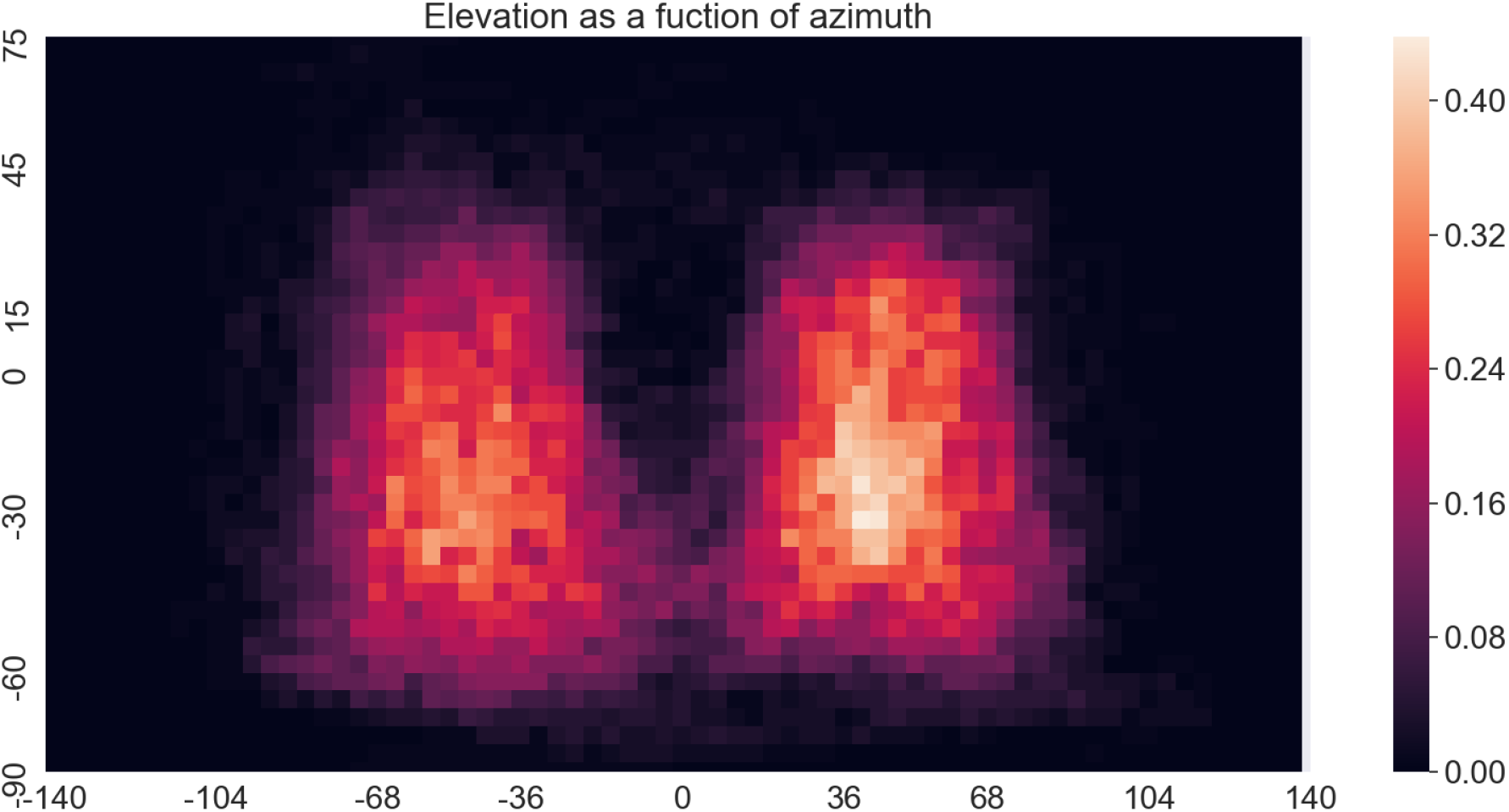
Probability density of the tip of the antenna passing by a specific region (a specific azimuth and elevation angle). Each pixel represents 4 degrees. The color of each pixel corresponds to the proportion of videos where the antenna tip of the insect passed at least once by the corresponding elevation and azimuth (121 videos).

To roughly evaluate the impact of the antennal motion on odor sampling, the volume sampled by each antenna was calculated. We then calculated the rate of sampling of the oscillating antenna after the insect reaction and compared it to the rate of sampling of a same size antenna immobile in the airflow (see example fig S4). We found a significative (one sample t-test N=121, p=10^−16^) increase in the sampling rate of the moving antenna, on average 1.2mm^3^/s. The base sampling rate of the static antenna in the airflow was 21.35 mm^3^/s. This meant that the antenna could double the volume of air sampled in 18 seconds of oscillation. No significative correlation was found between the diffusivity of the odor stimulus and the rate of sampling by the antenna (Spearman’s rank correlation N=121, p=0.097 and rho=0.15) (Fig S5).

### 3.2 Oscillations and their frequency components

Since the motion of the insect antennae presents slow and fast oscillations, we conducted a frequency analysis of the antennal motion of the insect after it received a stimulus. For this, the fast Fourier transformation of Scipy 1.6.1 was used with a Hanning window. It was necessary to detrend our oscillation data to center our signal on zero and remove any linear drift stemming from low frequency noise prior to frequency analysis. Because our videos were recorded at 300fps, we were able to do a spectral analysis up to 150Hz. The angular position of the antenna after the start of the reaction shows a maximum in amplitude between 0.5 and 2Hz (Fig 4). This peak is especially visible for the up and down motion of the antenna (the elevation angle).

**Fig 4:**
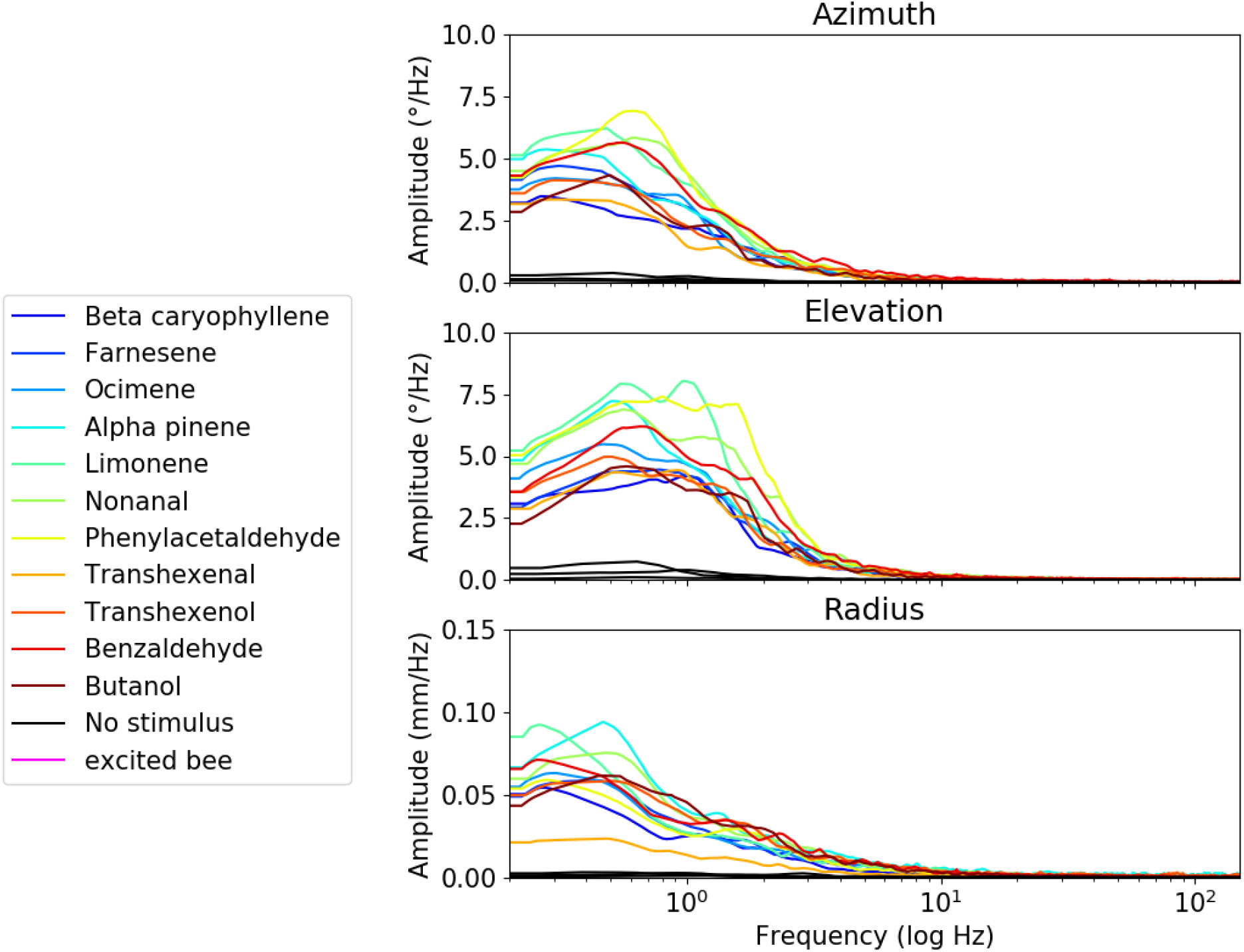
Spectral analysis of the motion of the tip of the antennae along each coordinate. Mean response over all bees for each odor (N=149).

The high frequency oscillations were not noticeable on the Fourier transform of the regular videos since their amplitude are too small. We thus recorded zoomed videos with higher frequencies. A 6^th^ order high pass Butterworth filter with a cutoff at 20 Hz was used to dampen the low frequencies of the motion. Seven trials on six insects were averaged. The Fourier transformation of the position of the tip of the antenna shows a spike between 40 and 60Hz in the amplitude of the azimuthal motion but not of the radial motion (Fig S6). Furthermore, the buzzing of the bumblebee (vibrations caused by the wing motion) was also accidentally recorded on two occasions while preparing the insect. This behavior was found to cause even higher frequencies of oscillations in the whole body, that propagated to the antennae. Those antennal vibrations were very visible and reached the frequency of the insect wingbeats (^70^), at around 200Hz (Fig S6).

### 3.3 Correlation of motions between the two antennae

Since the airflow on the antennae is coming from the right of the insect at 35mm/s and since the antennae are about 10mm apart, we could expect the right antenna to start reacting about 0.3 second before the left one. We found however no significative delay between the start of the reaction of both antennae, for any stimuli used (one sample t-test for each stimulus p>0.05 and one sample t-test for all stimulus N=121, p=0.9985) (Fig S7). It seems that bumblebees move both antennae together once an odor is sensed by any of the two antennae. No correlation was found either between the diffusivity of the odors and the delay of the antennae (Spearman’s rank correlation, N=121, p=0.77, rho= -0.027).

We also investigated an eventual dephasing of the motion of the antennae that would be caused by the lateralized stimulus. To do that, a sliding correlogram was realized by plotting the correlation coefficient of the two antennae motion for each angular coordinate with respect to the time lag between them (Fig S8). Since no prominent peak in correlation is visible, there is no constant dephasing between the antennae.

Finally, we analyzed the correlation between the motion of the antennae over time using a sliding window. A peak in the correlation of the antennae at the onset of the reaction was found. Specifically, the antennae move opposite in their left-right (azimuth) motion and move together in their up-down (elevation) motion. After the onset of the bumblebee reaction, this correlation disappears, and the antennae move independently (Fig 5).

**Fig 5:**
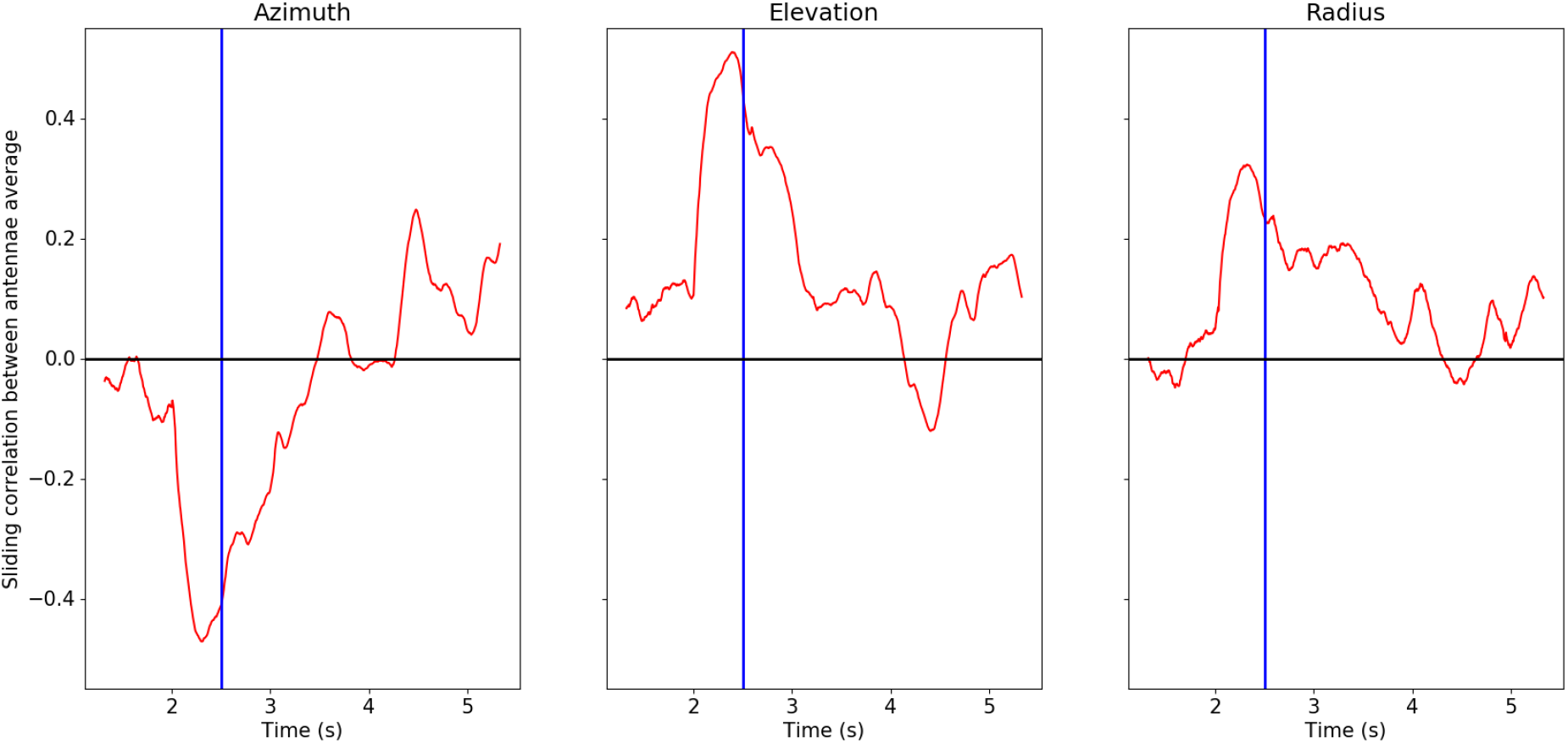
Sliding averaged correlation coefficient between the antennae for all stimulated bumblebees (N=121). The bees’ responses were synchronized with antennal deployment occurring at 2 seconds (blue bar).

### 3.4 Influence of the odor on the antennae motion

In order to reveal an influence of the odor sensed on the reaction of the insect, the amplitude and frequency of the motion for each odor were compared. No clear relationship between odor property and frequency of motion was found. The frequency of the motion stayed around 1Hz for the vertical oscillation and around 0.5Hz for the horizontal oscillations (Fig 4). An analysis of the amplitude of the motion of the antennae did not show an influence of the odor used either but demonstrated a difference between odor stimulation and control or excitation (Fig 6).

**Fig 6:**
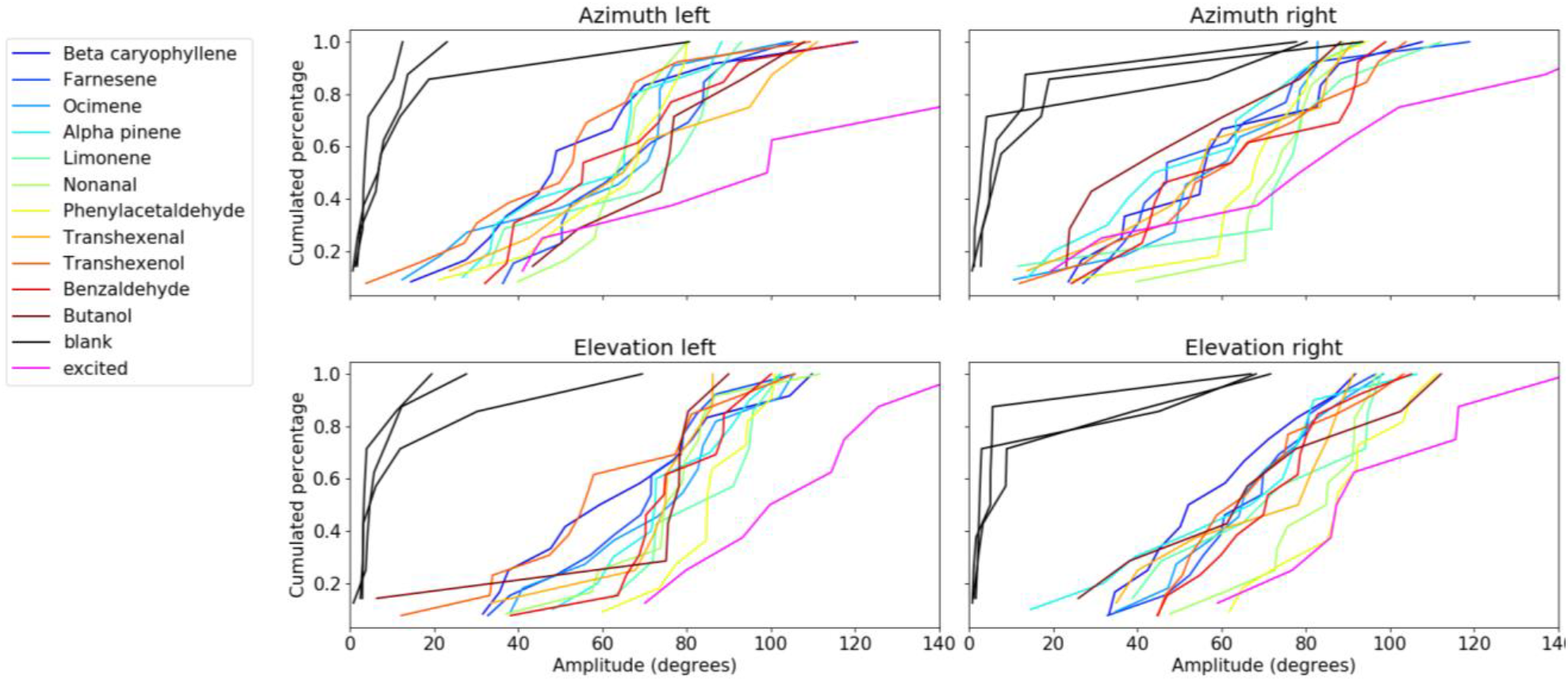
Cumulated percentages of bumblebees oscillating their antennae at a specific amplitude (N=149).

We also analyzed the maximum stroke speed of the insect antennae. We again found no clear differences between the stimuli used. The maximum stroke speed was always around 50 mm/s for all stimuli. The stroke speed was around 12mm/s for blank stimuli and 90mm/s for excited bumblebee (Fig 7).

**Fig 7:**
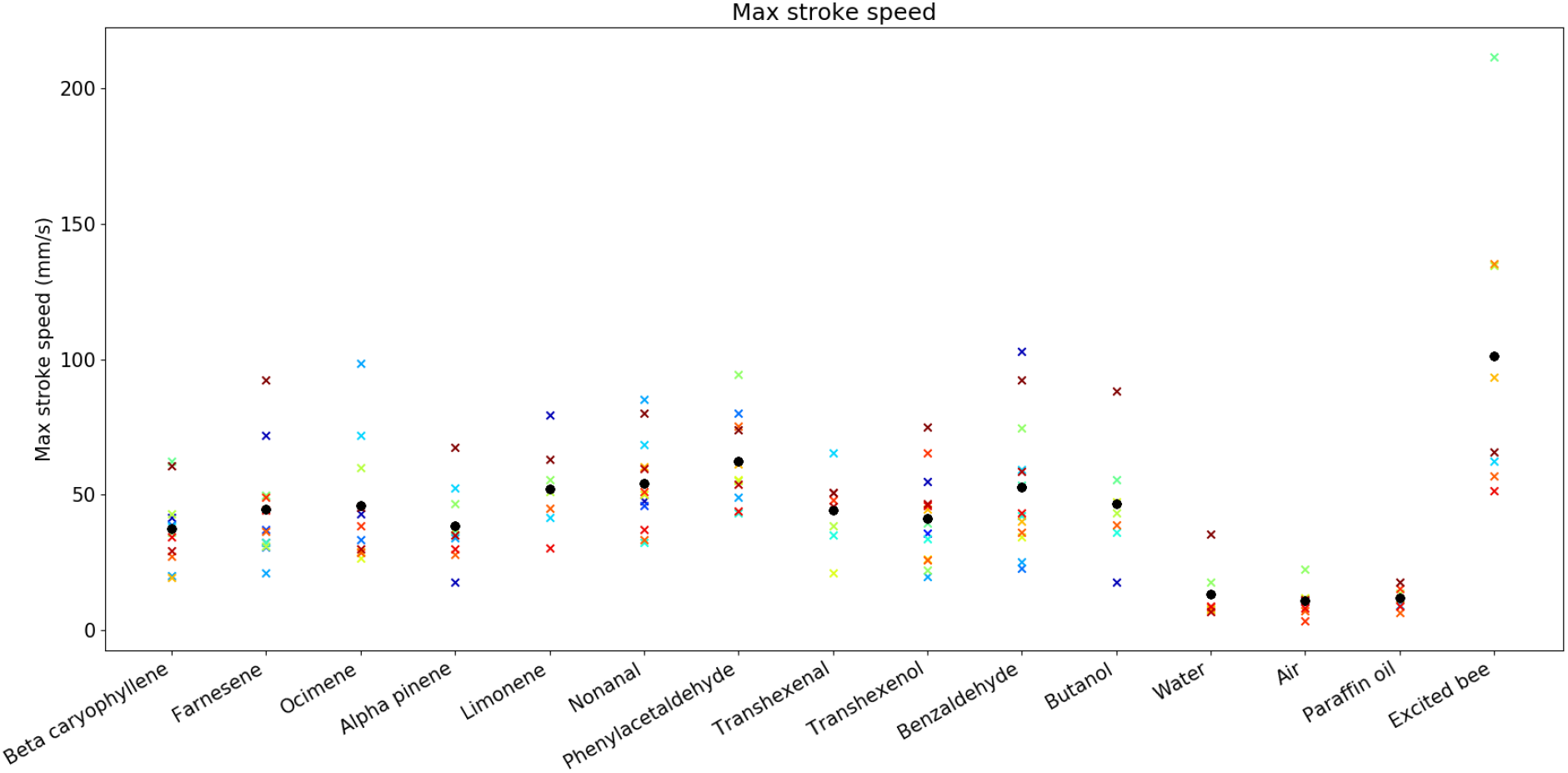
Maximum stroke speed of the tip of the bumblebee antennae for different stimuli. The black dot is the average of all bees for each stimulus, and each bee is represented by a color (N=149).

We finally looked for differences between onsets of the reaction and searched for correlation of these onset patterns with odor compounds detection. Again, no differences in the start of the motion due to the stimuli could be detected.

## 4 Discussion

In this study, we described and analyzed the active olfaction strategy of bumblebees. We show that bumblebees use a stereotyped antennal motion response when faced with an olfactory stimulus. This response is the combination of two oscillations of the antennae. The first is at high frequency and small amplitude and is constantly present regardless of the stimulation. The second is at low frequency, but with large amplitude, and is triggered by the odor stimulus. Both antennae reacted simultaneously, in a predefined pattern. We first discuss the limits of the experimental set-up of our study, then compare our results with those observed in previous works. We end the discussion by an interpretation of the movement pattern in terms of active search strategy.

### 4.1 Limits of the study

We made several decisions that impacted our results. First, we had to restrain the motion of the insect during filming; only its antennae were free to move. This setup partially restricted the movement of its head and could have modified its reaction somewhat as it has been shown that tethering bees affect their response to visual conditioning (^71^). Second, we introduced the odor stimuli from the right side of the insect, to be able to film its front side. This means that the odor reached the left antenna of the insect with a delay relatively to the right antenna. Since no lateralized response of the insect was observed, this did not seem to impact its reaction. Third, we used common odors as stimuli. The ratio of largest on smallest diffusivities and molecular weights was about two and a larger difference in those characteristics might have caused larger changes in the behavior observed. Odorant with more extreme diffusivity differences exist, Methanol (^72^) (16.3 mm^2^/s) and diterpenes like abietic acid (^73^) (4.24 mm^2^/s) are such compounds, but they are way less common in plant odors and have not been used for olfactory tasks on bees. To avoid causing unnatural reaction of the bees to these uncommon compounds we refrained from using them. We also used pure odor instead of cocktails of molecules such as those emitted by flowers for simplicity. This is a common decision in the research on bee olfaction (^53–55,57–60^). Fourth, our only way to identify the onset of the reactions is using an antennal motion threshold, since there was a delay for the odor to reach the insect after it was added to the continuous airflow. This limited our accuracy. Fifth, DEHS droplets were used to image the flow delivered to the insect. This showed that the airflow was laminar when it reached the insect but not homogeneously mixed. Fortunately, the patches of clean air were small with respect to the antennae. This ensured a systematic contact with the antennae, and the lack of complete odor mixing was not considered a problem. It is to be furthermore mentioned that the DEHS droplets used in the PIV measurement have a diffusion coefficient nearly 4 orders of magnitude lower than our less diffusive odor (supplementary material). As such, our stimuli would therefore diffuse more and be more homogeneous than what we observed (Fig S1). Finally, due to the length of the experiment, the weather condition of the experiments and the age of the hive changed between the first and the last insect testing. The temperature and hygrometry were variable between each test. Nevertheless, those differences did not seem to impact the insects’ stereotypical reactions. Also, because we did our experiments on any bumblebee exiting the hive, we do not know the age or the degree of experience of the bees we tested.

### 4.2 Comparison with former studies

Our study is the first to analyze the 3D motion of bee antennae, in contrast to the studies on the movements of the bees’ antennae caused by odor stimulation (^40–44^) done in 2D. Those studies were done on honeybees and showed that they move their antenna from front to back in an oscillating manner. Birgiolas et al (^44^) study recorded oscillations with an amplitude of 100 degrees and a frequency of 0.5 to 3 Hz, while Chole et al (^43^) recorded oscillations of the same magnitude but with the dominant frequencies below 0.5Hz. Erber et al (^40^) measured frequency of 0.5 Hz for resting bees and up to 1 Hz for bees stimulated by geraniol. Because we recorded the antenna motion in 3D, we were able to decompose the motion of the antenna into its vertical and horizontal oscillation. We found that the oscillation described in the 2D studies come mainly from the vertical oscillation of the antennae. The amplitude we measured for those oscillation was a bit lower than the one measured in the first two studies (Erber et al did not measure the oscillation amplitude). The amplitude of our oscillation was around 60 degrees for the horizontal oscillations and 70 degrees for the vertical oscillations. We did find around 100 degrees of amplitude of vertical oscillation in the excited bumblebees. We also found frequencies similar to the study by Birgiolas et al and Erber et al between 0.5 and 2 Hz. The high frequency oscillations of the antennae we measured were not mentioned by any other studies on bees probably due to the measurement precision required, but similar high frequencies (around 40 Hz) have been observed in moth antennae (^18^). They have been shown to create vortices that could help increase the capture of molecules on the antenna (^74^). A similar phenomenon could be the reason of the high frequency oscillations observed in the same condition in our insects, but further study is needed.

The speed of the strokes of *Apis mellifera* antennae, and the link between an increase in the speed of the air around the antenna caused by those strokes and EAG responses of the insect was investigated by Peteraitis (^41^). This study found antenna movement speed from 0 to 150mm/s, corresponding to the same range we observed with our bumblebees. The fastest stroke speed we observed were around 135mm/s, but they were exceptional and only happened for excited bumblebees. When stimulated by an odor, most bumblebees moved their antennae around 50mm/s. Lambin et al (^42^) studied the angular speed of the antenna when stimulated by an odor. They found that citral caused an increase in angular speed of the antenna in the 6 seconds after the start of the stimulation from 18°/s to 22°/s on average. We did find an increase in antenna angular speed in the seconds following the stimulus, but the angular speed observed were closer to 30°/s before and 150°/s after stimulation on average. The angular speed of the antennae recorded by Chole et al are also much lower than the speed we recorded or that any other study reported at around 6°/s.

Both Birgiolas et al and Chole et al described the initial reaction of the antennae to the odor as moving away from the source of the odor. Erber et al and Suzuki (^39^) on the contrary described the antennae as moving toward the odor source. We did not find a similar tendency. Instead, we observed both antennae deploying and raising up at the start of the stimulation. This could have been interpreted as moving away from a source of odor in Birgiolas et al and Chole et al studies. Chole et al also noted that the bee antenna moved closer to each other when stimulated by the odor before conditioning. We observed a similar reaction in the bumblebees tested. This is also observed in the anticorrelation of the azimuthal motion we found between antennae at the start of the reaction. We furthermore observed an increase in the distance between the tip and the base of the antenna at the start of the stimulation that Chole et al did not observe. An explanation for this discrepancy could be that bees were not calmed down before stimulation in that study. Their antennae were already deployed and moving. Our bumblebees were by contrast calm and their antennae immobile, if not folded, prior to the stimulation. Another explanation could be that Chole et al 2D measurement were not able to accurately measure the radial movement of the antennae tips. This could also be the explanation for the large variability in the radial position of the antennae that they measured in contrast to the very low radial motion we measured after the deployment of the antennae.

### 4.3 Odor sampling strategy

Bees displayed the same amount of variability of the antennae motions between odors as between individual insects despite the large variability of molecules. This could mean that, from a physical point of view, the diffusion coefficients of the odors in air are not different enough to require an adaptation of the antennal motion to each substance. The same type of antennal search strategy would then work for the large variability of molecules a bee would encounter.

The vertical oscillations could be used simply to increase the airflow on the antenna and hence the capture of the molecules. They would then serve a similar purpose as sniffing in vertebrate, increasing the airflow on the sensory surfaces and preventing the adaptation of the sensory neuron to the stimuli (^4^). The horizontal movement would by contrast serve to sample the air in specific regions around the insect to search for the source of odor by selecting the plane of the vertical oscillation. This type of strategy can be observed in the casting behavior of other animals (^5,13^) and was described by Chen et al (^75^) as energy-constrained proportional betting. It serves to maximize information gain during sampling. Oscillating the antenna in any direction would also allow the insect to sample an area much larger than the surface of its antenna. This could prove useful in the more natural situation of heterogeneously mixed air, to increase the chances of encountering a plume of odor. Further studies investigating those hypotheses would need to record the 3D motion of the antennae concurrently with a precise mapping of the turbulent flow and chemical plumes, very much in the spirit of those conducted by Koehl et al on crayfishes and followers (^22,76,77^).

