## Supplementary file for "Active sensing in bees through antennal movements is independent of odor molecule"

### 1 Diffusion of DEHS

Using a similar method to the analysis of water droplet diffusion for virus propagation ([1]) we analyzed the diffusion speed of the DEHS droplet used to emulate an odor stimulus.

The diffusion coefficient of a particle in air can be calculated at the level of binary collision of air molecule with the particle if its size is comparable or smaller than the mean free path in air (around 100nm):

(1)

$$D_1 = \frac{3(k_B T)^{3/2}}{2^{3/2}\pi^{1/2}m^{1/2}PR^2}$$

Where $k_B$is the Boltzmann constant, T the temperature in Kelvin, m the average mass of the molecules, P the pressure and R the particles radius.

If the particle is larger, its diffusion coefficient can be calculated using fluid dynamic:

(2)

$$D_2 = \frac{k_B T}{6\pi\eta R}$$

Where $\eta$ is the dynamic air viscosity ($1.85*10^{-5}$ kg/(m*s) at 25°C).

For a given particle radius, the diffusion coefficient should be the maximum between D1 and D2. In our case, the DEHS particles used measure 0.2µm, giving a radius of 100nm. The equation gives D1=$2.28*10^{-11}$ $m^2/s$ and D2=$1.19*10^{-10}$ $m^2/s$. An approximation of the diffusion coefficient of our droplet should then be $1.19*10^{-10}$ $m^2/s$, which is way larger than our less diffusive odor, farnesene, with a diffusion coefficient of $4.91*10^{-6}$ $m^2/s$.

**Supplementary figures**

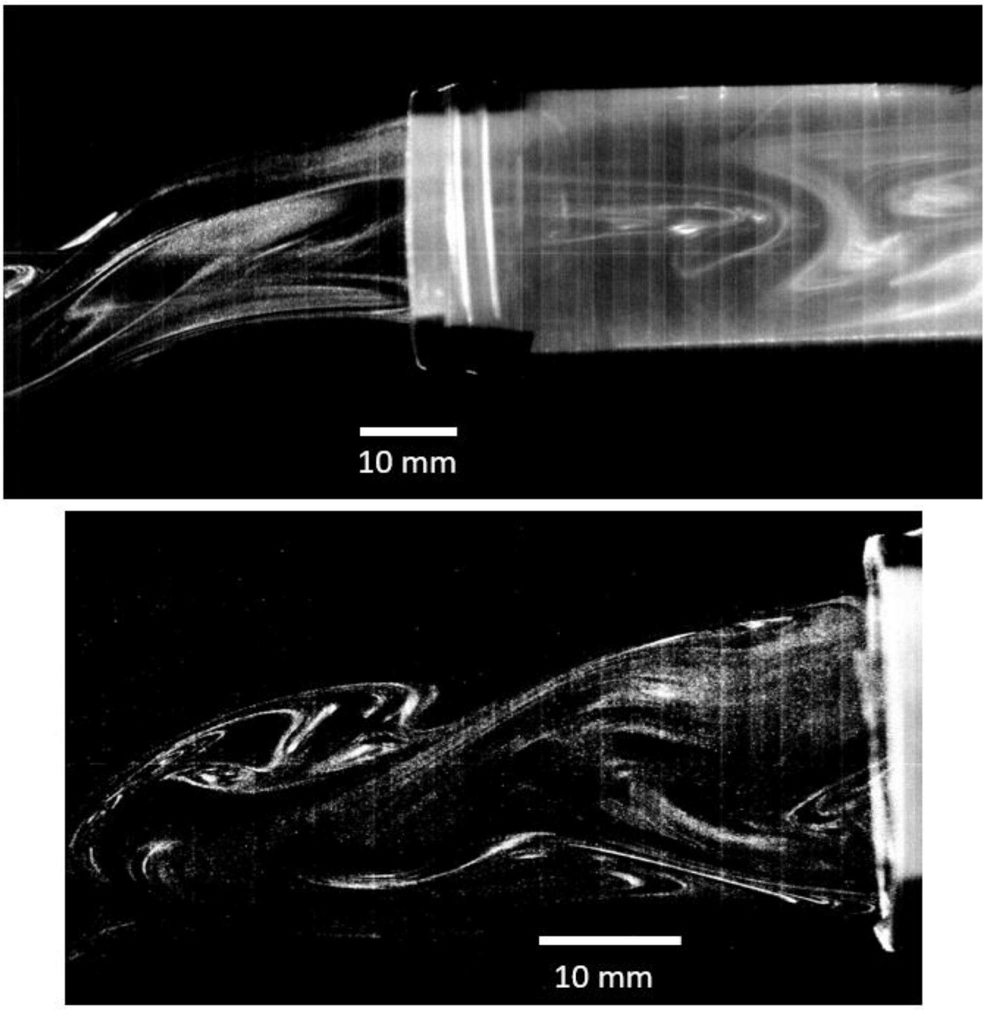


Fig S1: Pictures of the distribution of DEHS droplets inserted in the delivering tube in the same way the odor stimulus was. The heterogeneous mixing is quite visible.

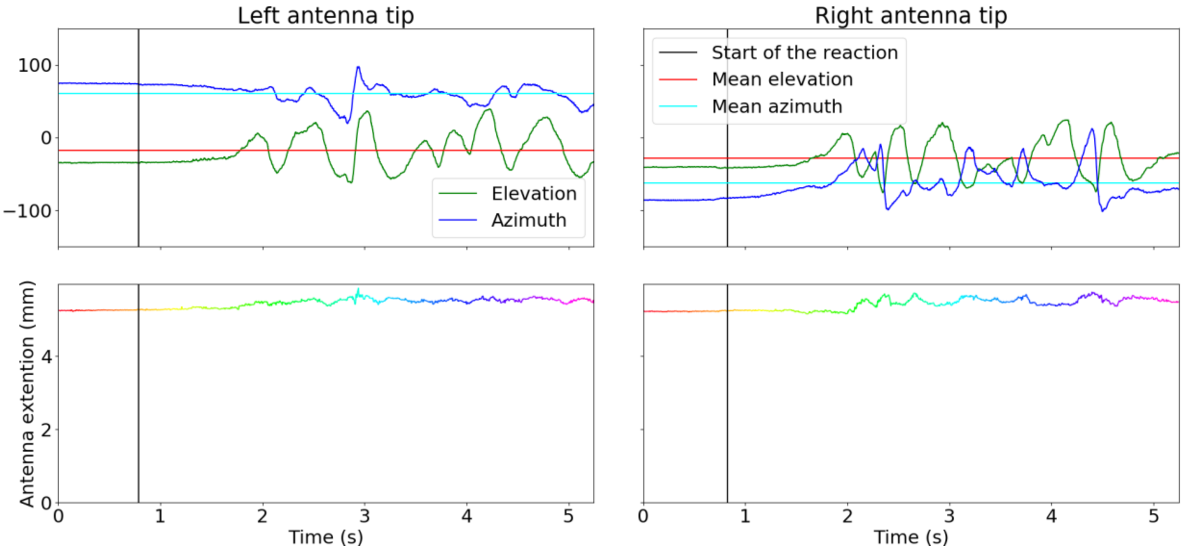


Fig S2: Typical reaction of a bumblebee to a stimulation. At the top, the angular position of the tip of each antenna. At the bottom, the radial position of the tip of each antenna. The scale of the radial position highlights the small range of radial motion compared to the length of the antennae.

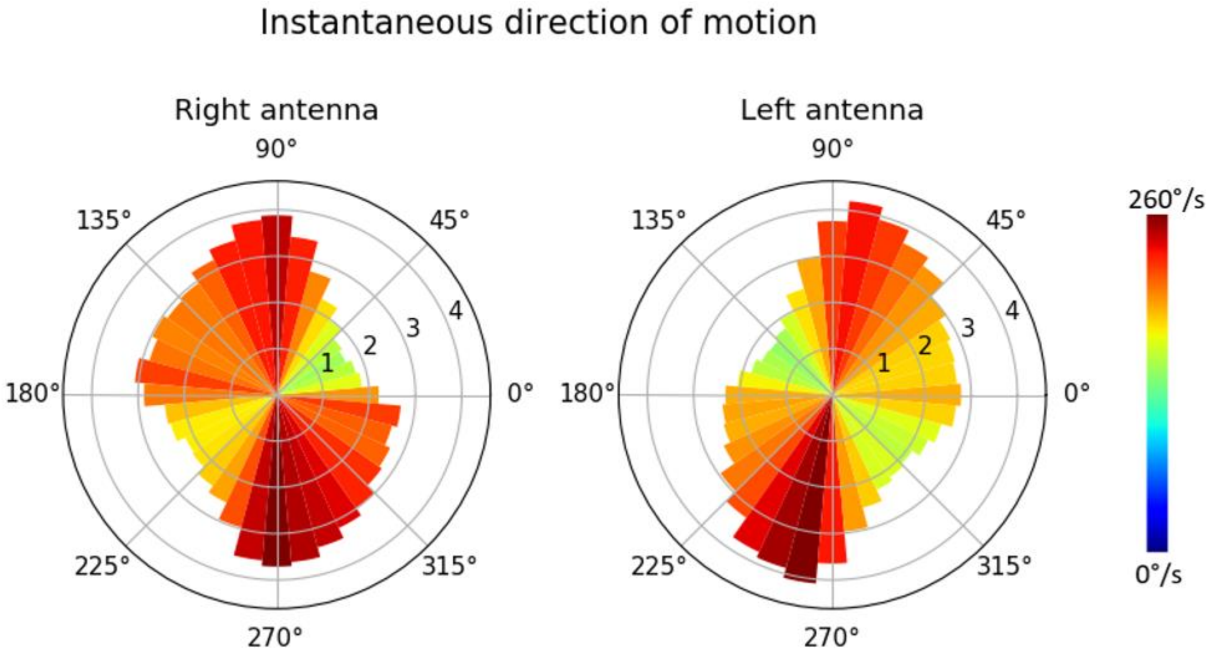


Fig S3: Histogram of the instantaneous direction of motion of the tip on the antenna during the insect reaction as well as the motion speed in theses directions. The length of the bar represents the percentages of frames with the antenna tip moving in this direction. The color of the bar represents the median speed in a specific direction.

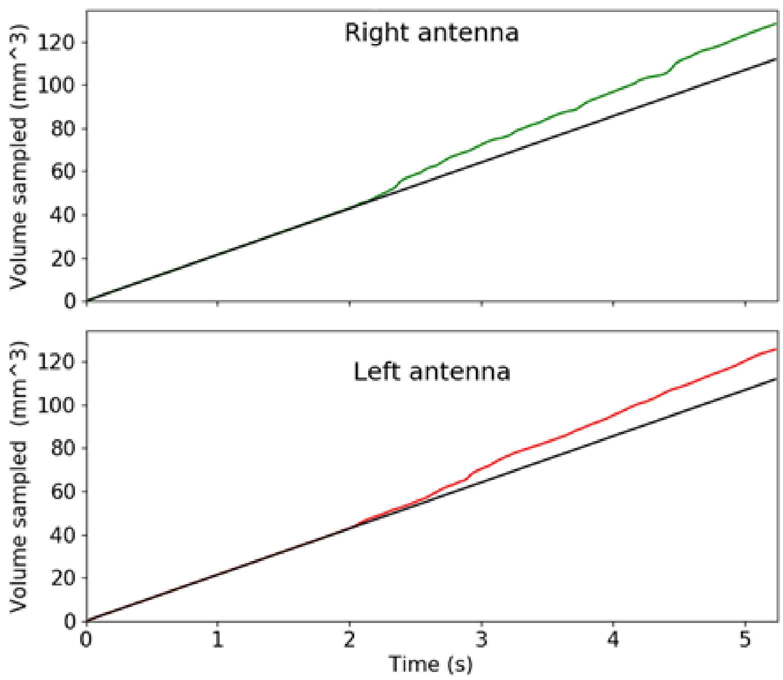


Fig S4: Example of the volume sampled by the insect antennae during its motion (in color) compared to the volume sampled by a static antenna in the same airflow (black).

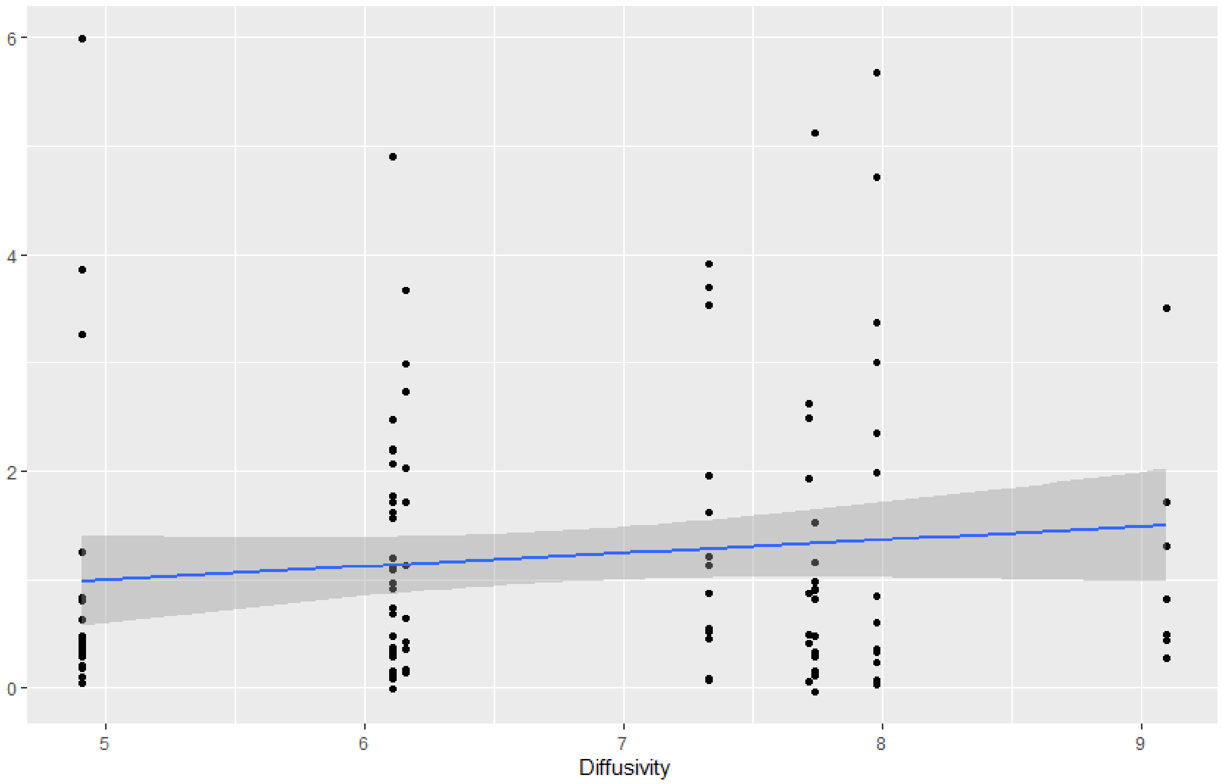


Fig S5: Air sampling rate of the antenna with respect to the diffusivity of the stimulus molecule. The linear regression and the 95% confidence interval are also shown.

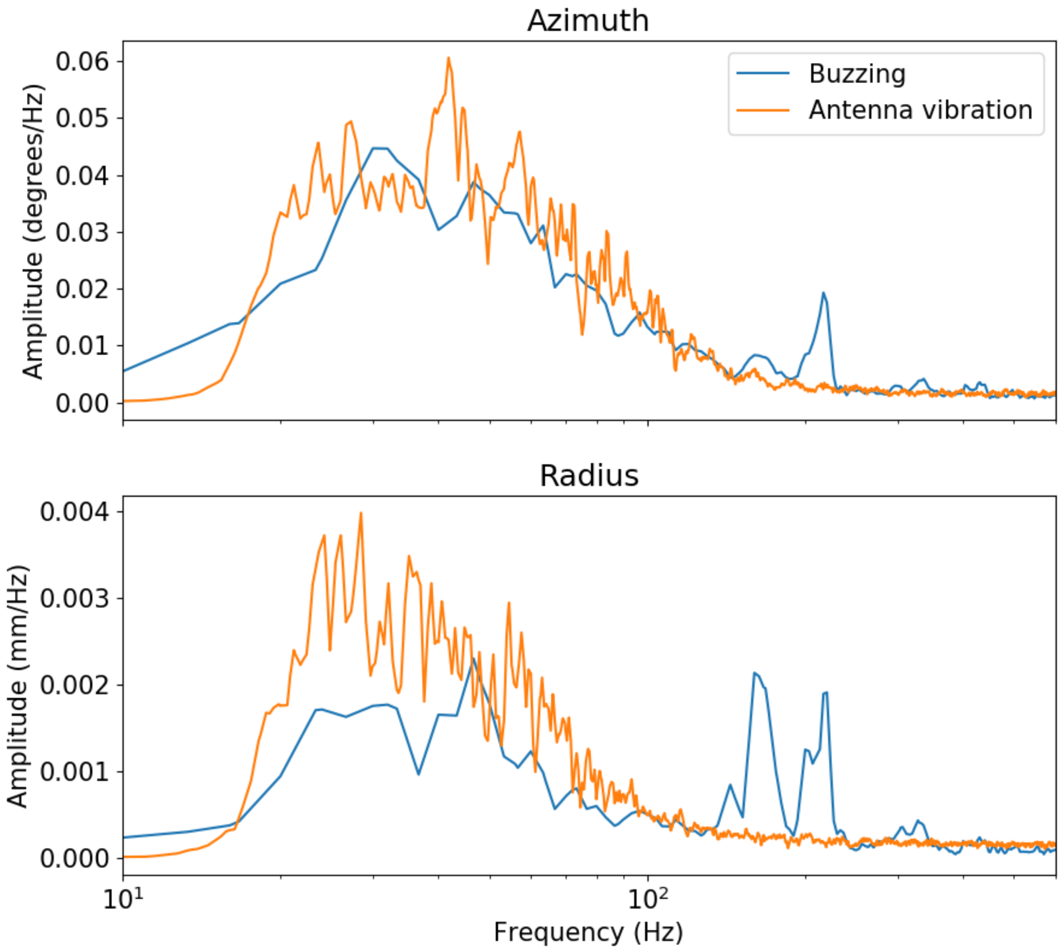


Fig S6: Spectral analysis of the motion of the tip of the antennae along each coordinate for the high-speed recordings (angular and radial position). Comparison of the high frequency oscillation with the buzzing of the insect. The frequency scale is logarithmic.

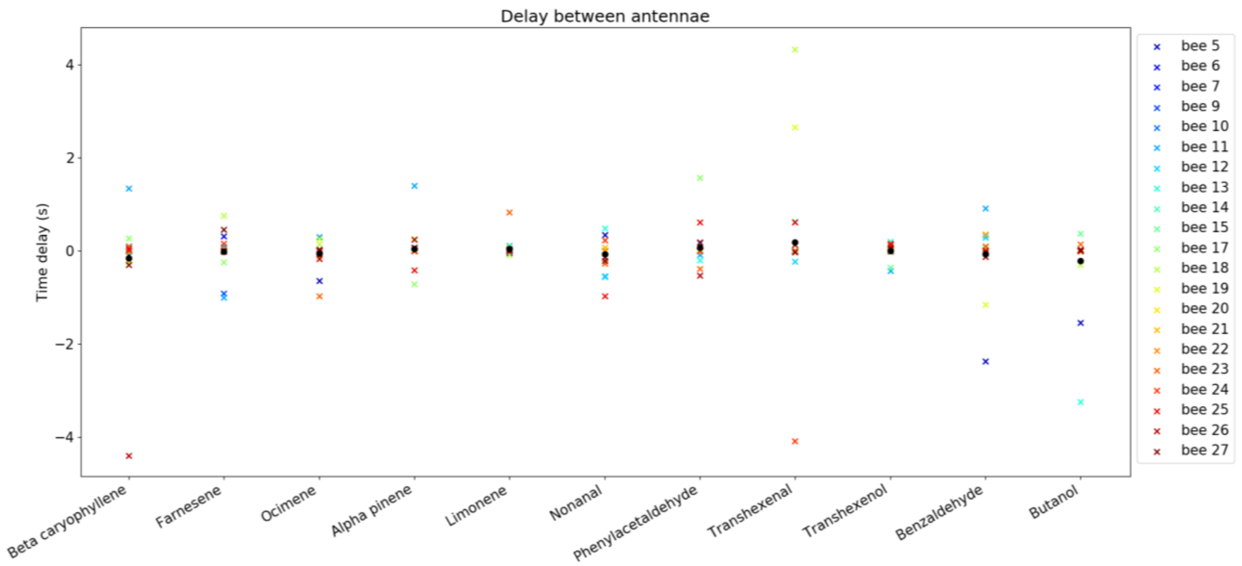


Fig S7: Delay (seconds) between the antennae of the insect depending on the stimuli. The black dot is the average. A negative delay mean that the left antenna reacts first.

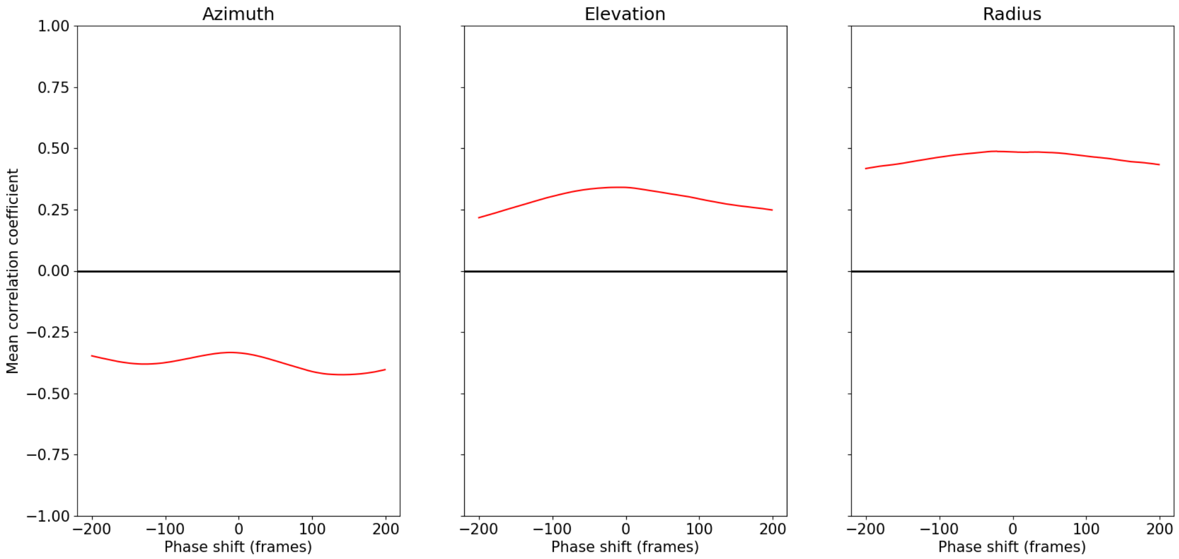


Fig S8: Correlation coefficient of the antennae for each coordinate with respect to the time lag between them (in second)
